# Statistical Inference Relief (STIR) feature selection

**DOI:** 10.1101/359224

**Authors:** Trang T. Le, Ryan J. Urbanowicz, Jason H. Moore, Brett A. McKinney

**Affiliations:** 1nstitute for Biomedical Informatics, University of Pennsylvania, Philadelphia, PA 19104, USA; Department of Mathematics, University of Tulsa, Tulsa, OK 74104, USA; Tandy School of Computer Science, University of Tulsa, Tulsa, OK 74104, USA

## Abstract

**Motivation:** Relief is a family of machine learning algorithms that uses nearest-neighbors to select features whose association with an outcome may be due to epistasis or statistical interactions with other features in high-dimensional data. Relief-based estimators are non-parametric in the statistical sense that they do not have a parameterized model with an underlying probability distribution for the estimator, making it difficult to determine the statistical significance of Relief-based attribute estimates. Thus, a statistical inferential formalism is needed to avoid imposing arbitrary thresholds to select the most important features.

**Methods:** We reconceptualize the Relief-based feature selection algorithm to create a new family of STatistical Inference Relief (STIR) estimators that retains the ability to identify interactions while incorporating sample variance of the nearest neighbor distances into the attribute importance estimation. This variance permits the calculation of statistical significance of features and adjustment for multiple testing of Relief-based scores. Specifically, we develop a pseudo t-test version of Relief-based algorithms for case-control data.

**Results:** We demonstrate the statistical power and control of type I error of the STIR family of feature selection methods on a panel of simulated data that exhibits properties reflected in real gene expression data, including main effects and network interaction effects. We compare the performance of STIR when the adaptive radius method is used as the nearest neighbor constructor with STIR when the fixed-*k* nearest neighbor constructor is used. We apply STIR to real RNA-Seq data from a study of major depressive disorder and discuss STIR’s straightforward extension to genome-wide association studies.

**Availability:** Code and data available at http://insilico.utulsa.edu/software/STIR.

## 1 Introduction

Epistasis is a well known concept in genetics that can be statistically modeled as a deviation from the additive effect of DNA variants on a phenotype or trait. A similar effect can be observed at the gene expression level, where the phenotypic effect of one gene is modified depending on the expression of another gene (Park and Lehner (2013)). A manifestation of this “expression-epistasis” effect is differential co-expression (Lareau *et al*. (2015)). The embedding of these interactions in a regulatory network may lead to, not only pairwise interactions, but also higher-order epistasis network effects. Thus, feature selection methods are needed for highdimensional data – such as genome-wide association and gene expression studies – that are able to identify relevant features when their effect on a phenotype may be obscured by a complex interaction architecture.

Relief-based feature selection methods are known for their ability to identify interactions with computational efficiency based on nearest neighbor calculations in the high-dimensional feature space (Urbanowicz *et al*. (2018b); Kononenko *et al*. (1997); McKinney *et al*. (2009); Kira and Rendell (1992)). The early Relief-based algorithms used arbitrary parameter choices for the number of nearest neighbors and heuristic Relief-score thresholds for selecting the most important features. Recent work has been done to address the selection of the number of nearest neighbors, such as the constant neighborhood radius in spatially uniform ReliefF (SURF) (Greene *et al*. (2009)), adaptive radii in multiSURF (Urbanowicz *et al*. (2018a)) and feature-specific optimal *k* in ReliefSeq (McKinney *et al*. (2013)). However, until the current study, the threshold for selecting the top predictors has remained arbitrary because Relief scores have not had a null distribution.

Methods like ANOVA and the generalized linear model have parametric probability distribution assumptions that easily and efficiently permit the calculation of p-values. However, these methods are not able to detect interactions unless each interaction term is explicitly included in the model. Explicit interaction modeling becomes computationally intractable for high-dimensional data and/or higher-order interactions due to the combinatorial explosion of hypothesis tests. Meanwhile, Relief ranks the importance of each attribute separately, like a univariate method, but its ranking accounts for dependencies between all other attributes, making it “omnivariate.” Next we discuss the mechanism that Relief-based methods use to incorporate interaction effects in importance scores while circumventing the combinatorial explosion.

When updating a target attribute’s importance score for an instance in the data, Relief accounts for variation between all other attributes by using nearest neighbors of the instance, as computed in the space of all attributes. For the target attribute, a hyper-dimensional decision boundary *(i.e*., in the space of all attributes) is computed for each instance, and the attribute’s score is updated from the neighbors near this boundary. In effect, Relief creates a high-dimensional nonlinear decision boundary localized at each instance to discriminate between its nearest hits (same class) and misses (different class). Pairwise attribute interactions are not explicitly calculated in Relief, but pairs of attributes that interact conditionally on the outcome variable will both have similarly high Relief importance scores. Relief also has the ability to identify higher order interactions, again without explicit calculation of *n*-way interactions.

Relief-based methods are, thus, an excellent tool for detecting interactions, but, as noted, there remains the challenge of determining statistical thresholds or statistical significance. With the aim of addressing this challenge, we recently developed a mixture model and a permutation approach to estimate statistical thresholds for ReliefF and network centrality scores (Lareau *et al*. (2015)). However, permutation testing can be computationally prohibitive. To address this issue, in the current study we introduce a new family of Relief-based algorithms that allows for statistical inference and false discovery rate adjustment.

The new STatistical Inference Relief (STIR) formalism represents a new type of Relief-based score that follows a pseudo t-distribution. In a precursor of STIR, we recently demonstrated that scores from the standard Relief algorithm are equivalent to a difference of mean attribute value differences between nearest hit and miss groups (McKinney *et al*. (2013)). This equivalence suggests a reformulation of Relief scores that accounts for the variance within and between groups. STIR in the current study is able to detect attributes whose association with the phenotype may be due to higher-order interactions while simultaneously assigning statistical significance to the attribute scores. The STIR formalism applies to the broad family of Relief-based algorithms, including Relief with fixed *k* and multiSURF.

The paper is organized as follows. In the Methods section, we develop the new formalism of STIR that enables the calculation of the STIR pseudo t-statistics (STIR scores) and statistical significance of these scores. We discuss our simulation strategy involving main effects and realistic network interaction effects of varying strengths, sample sizes, and number of attributes. In the Results section, we apply the STIR method to the panel of simulated data to assess power and false discovery rates. We use STIR to obtain FDR-adjusted statistical significance levels and compare with permutation testing. We compare STIR using *k* neighbors (constant for each instance) with multiSURF (variable for each instance) as the Relief-based nearest-neighbor algorithms. We apply STIR to a real RNA-Seq dataset from a study of major depressive disorder, and we note that STIR also applies to GWAS data. In the Conclusion section, we discuss challenges and opportunities for further development of the new STIR family of feature selection algorithms.

## 2 Materials and Methods

In this section, we develop the mathematical formalism for computing the statistical significance of Relief-based scores for feature selection for binary-class (case-control) data. We generalize the STIR formalism to all current nearest-neighbor methods, discuss the relationship between multiSURF and fixed-*k* methods, and demonstrate how the reformulation of Relief-based algorithms can be used to improve the computational performance of the algorithms.

### 2.1 Reformulation of Relief-based estimators

#### 2.1.1 Diff function and nearest neighbors

Before importance scores can be computed for each attribute, Relief-based algorithms identify the nearest neighbors in the space all attributes. The distance between instances *R_i_* and *R_j_* is calculated in the space of all attributes *a* ∈ *A*, typically using a Manhattan (*q* = 1) metric but may also use a Euclidean (*q* = 2) metric:

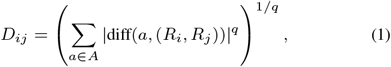

where the standard “diff” function between two instances *R_i_* and *R_j_* for a real-valued attribute *a* is:

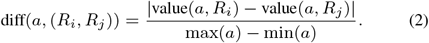

This diff is appropriate for gene expression and other real-valued predictors. For genome-wide association study (GWAS) data, where attributes are categorical, one simply modifies the diff, but the algorithm is otherwise unchanged. The diff function is part of the metric used by Relief methods to compute the distance matrix for finding nearest hit and miss neighbors, but the diff is also essential for computing the Relief importance scores, as will be seen in Sec. 2.1.3.

#### 2.1.2 Hit and miss nearest-neighbor ordered pairs

For a given instance *R_i_* (*i* ∈ 1,…, *m*), a hit is defined as a neighbor instance that has the same class label as that of *R_i_*, and a miss is a neighbor instance with a different class label from *R_i_*. In general Relief-based algorithms, one may represent the set of ordered pairs (*R_i_*, *M_j_i__* (*R_i_*)), or simply (*R_i_*, *M_j_i__*), of *m* instances *R_i_* with their nearest *k_M_i__* misses, *M_j_i__*, as nested sets:

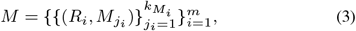

where the index *j_i_* for the inner set ranges from 1 to *k_M_i__*, which is the number of nearest miss neighbors for subject *R_i_*. The outer set ranges over all *m* instances. Similarly for hits, the set of ordered pairs (*R_i_*, *H_j_i__* (*R_i_*)) of *m* instances *R_i_* (*i* = 1,…, *m*) with their nearest hits, *H_j_i__*, may be written as

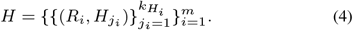

Note that in both miss and hit sets, the inner index *j_i_* depends on the outer index *i*. This is important for multiSURF, where each instance *R_i_* will, in general, have a different number of misses and hits (*k_M_i__* and *k_H_i__*) and these values may differ between instances. Thus, for multiSURF, the sets *M* and *H* can be thought of as irregular or ragged matrices of ordered pairs. For ReliefF algorithms, where the number of neighbors is constant across subjects, the hit and miss matrices are proper (non-ragged) matrices of ordered pairs.

#### 2.1.3 Reformulation of Relief-based estimators as difference of hit and miss means

Once the hit and miss groups, *H* (Eq. 4) and *M* (Eq. 3), are determined by the distance matrix *D_ij_* (Eq. 1) coupled with a neighborhood definition (*e.g*., ReliefF fixed number of neighbors *k* or multiSURF instance-dependent radius), we can compute average hit and miss diff means and attribute importance weights. We showed in Ref. (McKinney *et al*. (2013)) that the ReliefF importance weight for an attribute, *a*, can be expressed as a difference of mean diffs between hit and miss groups. Here we extend this difference to any Relief-based neighborhood scheme.

The mean diff for attribute *a* averaged over of all pairs of nearest-neighbor misses *M* (Eq. 3) can be expressed as

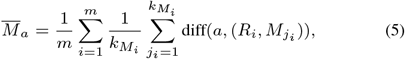

where *M_j_i__* is the *j*^th^ nearest neighbor from different classes of the *i*^th^ instance, *R_i_*, and *k_M_i__* is the number of nearest miss neighbors of instance *R_i_*. This scaling by 1/*k_M_i__* inside the sum makes the neighborhood average weighting consistent with multiSURF and with uniform neighborhood methods like SURF and ReliefF. For nearest neighbor hits, the mean is

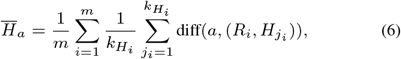

where, similarly, *k_H_i__* is the number of nearest hit neighbors of instance *R_i_*. The Relief-based importance score can then be expressed simply as

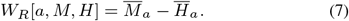

The formulation as a difference applies to any Relief-based algorithm. We will use Eq. (7) as the basis for computing permutation p-values for comparison purposes. However, as noted, permutation can have prohibitive computational times. Thus, in Sec. 2.2, we extend Eq. (7) to develop a Relief-based pseudo t-test and a more computationally efficient means of computing statistical significance of attributes.

#### 2.1.4 Performance optimization with the reformulation and ReliefF limits of general formalism

In our implementation of STIR on `R` ver. 3.4.4, we reshape all |*M*| and |*H*| ordered miss and hit pairs, *M* (Eq. 3) and *H* (Eq. 4), into |*M*| × 2 and |*H*| × 2 matrices to take advantage of `R`’s fast vectorization capability (Fig. 1). The reformulated algorithm may be optimized by pre-computing the neighborhood matrices *H* and *M* (Algorithm 2, line 7) and vectorizing the diff function so that we can simply perform vector subtraction (Algorithm 2, lines 11-13) and bypass the two nested `for` loops in the original algorithm (Algorithm 1, lines 8-11) in the calculation of the weight for each attribute. The description of the reformulated algorithm is simplified and allows for vectorization, which has a performance advantage over `for` loops in `R`.

**Fig. 1.**
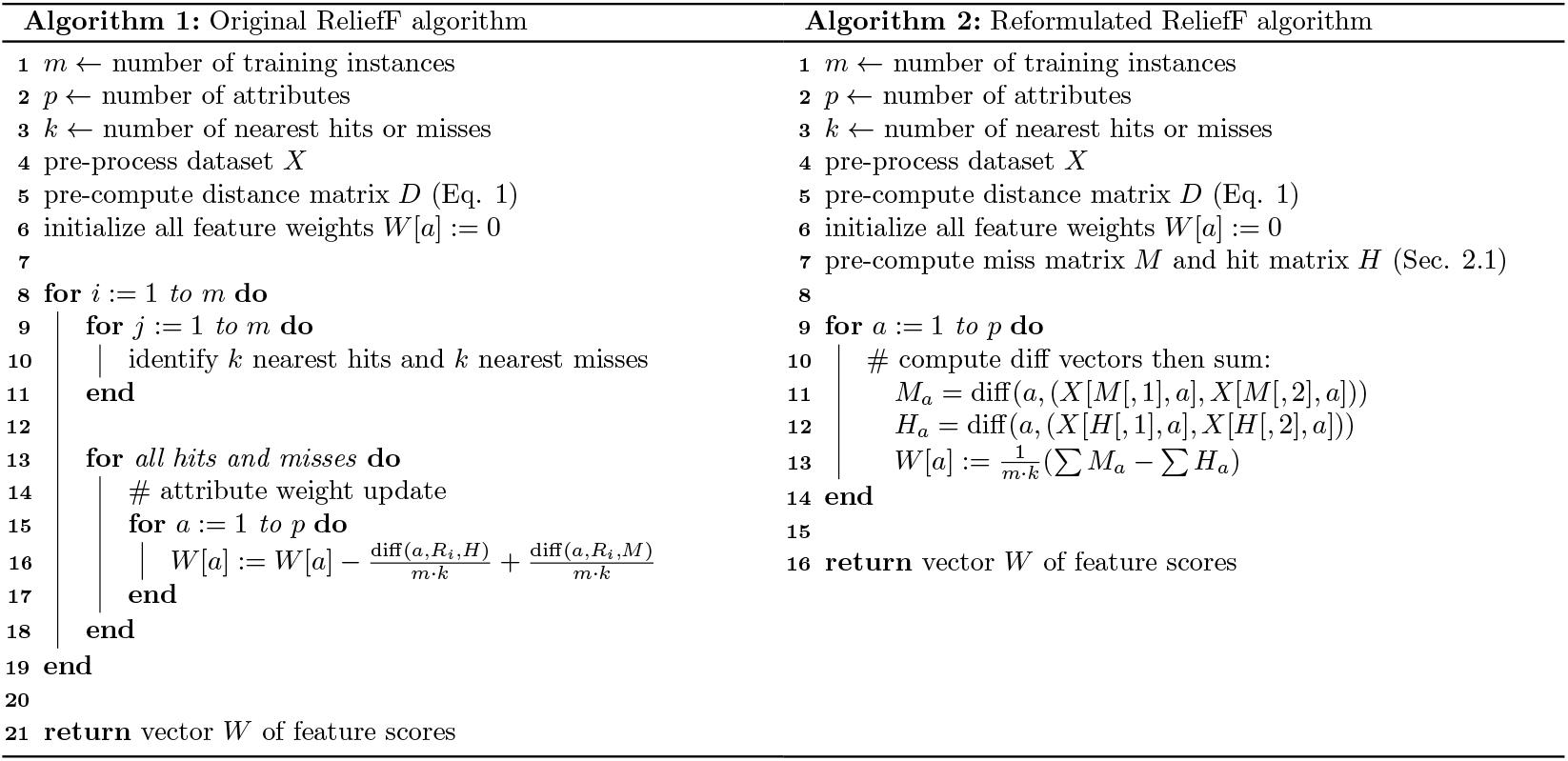
Comparison of the pseudo-code of the original ReliefF algorithm as implemented in ReBATE (Urbanowicz et al. (2018a)) (Algorithm 1, left) versus the reformulated version of ReliefF (Algorithm 2, right, based on Eq. 7 – line 13). The reformulated version allows for algorithm optimization by precomputing miss and hit matrices (Algorithm 2, line 7 – Sec. 2.1.4) and using a vectorized diff function (Algorithm 2, lines 11 and 12). The sums in line 13 are over all elements of *H_a_* and *M_a_* (all pairs of neighbors for all instances). The pseudo-code for STIR (Eq. 10) works similarly.

In the case of Relief-based methods with constant *k* (ReliefF), we have *k_M_i__* = *k_H_i__* = *k* ∀*i*, and Eqs. (5) and (6) become

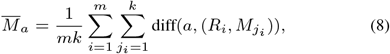

and

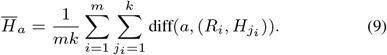

The ReliefF version of the reformualted score *W_R_* (Eq. 7) then follows directly.

### 2.2 Beyond Relief-based estimators: STatistical Inference for Relief (STIR)

We now introduce a new type of Relief-based score that incorporates the pooled standard deviations about the mean hit and miss diffs to transform the Relief-based score (*W_R_*) into a pseudo t-statistic, *W*_STIR_. For attribute *a*, we construct the following STIR weight (or STIR score) from the Relief difference of means (*W_R_* in Eq. 7) in the numerator and the standard error in the denominator:

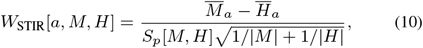

where 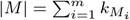 and 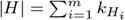 are the total number of miss and hit neighbors across all instances. The pooled standard deviation is

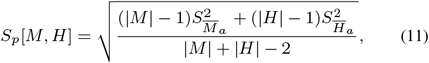

and the group variances are

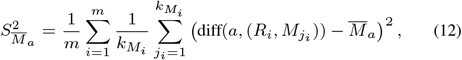

and

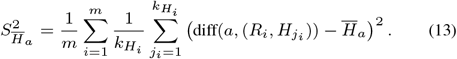

The pooled standard deviation above allows for unequal variances in the hit and miss nearest neighbor diffs and allows for a different number of diffs in the hit and miss groups, which is common for multiSURF. For Relief with fixed neighbors k, the above equations can be simplified by letting *k_M_i__* = *k_H_i__* = *k* and |*M*| = |*H*| = *mk*. The *W*_STIR_ score (Eq. 10) approximately follows a t-distribution from which we compute p-values. We use *df* = |*M*| + |*H*| — 2 as the degrees of freedom for calculating the p-value.

We highlight that STIR applies to any Relief-based algorithm. In this work, we focus on two different approaches for the neighbor finding algorithm (ReliefF and multiSURF) for use in STIR. ReliefF requires the user to specify a fixed *k* while multiSURF uses a neighborhood radius that varies for each instance (Urbanowicz *et al*. (2018a)). In multiSURF, the radius for each instance is the average of all distances of the instance to all other instances subtracted by half of their standard deviation. The multiSURF method counts another instance as a neighbor if it is within this radius. We show empirically for balanced datasets that a good constant-*k* approximation to the expected number of neighbors within the multiSURF radii is *k* = *m*/6. We show that the performance of STIR_*k*=*m*/6_ closely follows that of STIR-multiSURF.

### 2.3 Datasets and performance metrics

#### 2.3.1 Simulation methods

To address power and false positive performance of STIR, we use the simulation tool from our private Evaporative Cooling (pEC) software (Le *et al*. (2017)). This tool was designed to simulate realistic main effects, correlations, and interactions that one would expect in gene expression or resting-state fMRI data. In the current study, we first simulate main effect data with *m* = 100 subjects (50 cases and 50 controls) and *p* = 1000 real-valued attributes with 10% functional (true positive association with outcome). We chose a sample size consistent with real gene expression data but on the smaller end to demonstrate a more challenging scenario. Similarly, an effect size bias of *b* = 0.8 was selected to be sufficiently challenging with power approximately 40% (Le *et al*. (2017)). More details on the theoretical relationship between power and the simulation parameters is provided in Ref. (Le *et al*. (2017)).

One of the main advantages of Relief-based methods is the ability to detect statistical interactions. Thus, our second type of simulation uses the differential co-expression network-based simulation tool in pEC to simulate interactions. Full details of the simulation approach can be found in Refs. (Le *et al*. (2017); Lareau *et al*. (2015)). Briefly, we simulate *m* = 100 samples and *p* = 1000 attributes with 10% targeted for interaction. Starting with a dataset of random normal expression levels, we induce a coexpression network with Erdős-Rényi connectivity by making connected genes (*e.g*., *g_i_* and *g_j_*) have a linear dependence (*g_j_* = *g_i_* + *s*_int_) with average correlation noise *s*_int_. A *lower* value of *s*_int_ yields *higher* average co-expression and thus *higher* average interaction effect size.

The interaction is enforced by randomly targeting 10% of the attributes and permuting their values within the group of instances designated as cases. By permuting the values of the gene in cases, no main effect is created but the co-expression between the gene’s connections is destroyed in the case group, creating differential co-expression or interaction effects with that gene’s connections. We chose the 10% of targets randomly, which means that a few attributes may not have correlation with other attributes and hence may not actually be functional. On the other hand, other target attributes may be highly interconnected and, hence, may be involved in high-order interactions. This complexity of interactions and correlations makes assessing true/false positives/negatives challenging; however, our goal is to simulate realistic data and the 10% of targets is a reasonable surrogate for true associations. We use a relatively challenging interaction effect size *s*_int_ = 0.4. See Ref. (Le *et al*. (2017)) for further discussion of main effect and interaction effect sizes.

#### 2.3.2 Performance metrics

We compare the performance of STIR across Relief-based methods, with Relief permutation test, and with univariate t-test for both main and interaction effect simulations. We choose a univariate t-test as a comparison method for main effect simulations because it gauges the effect size and the t-test is an effective standard approach for detecting differential expression without multiple conditions or covariates. Specifically, a simulated main effect attribute is considered functional if its mean expression is significantly different between the two outcome groups. Moreover, the STIR p-values are analogous to a t-test. STIR p-values are computed from a t-test distribution from each attribute’s STIR score (Eq. 10). Relief-based permutation p-values are computed based on the reformulated Relief-based score (Eq. 7). For permutation, we first compute the observed score for each attribute. We then permute the class label 10,000 times, recomputing attribute scores for each permuted dataset. The fraction of permutations for which the observed score exceeds the permuted score is the attribute’s p-value.

All resulting p-values (STIR, permutation, and univariate t-test) are adjusted for multiple testing using the Benjamini-Hochberg procedure (Benjamini *et al*. (2001)). Attributes with adjusted p-values less than 0.05 are counted as a positive test (null hypothesis rejected), else the test is negative. We assess the performance of each method by averaging the following performance metrics across 100 replicates of each simulation scenario: True Negative Rate (TNR), Precision, and Recall of the statistical tests. We remind the reader of the following definitions applied for the detected attributes

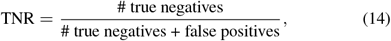

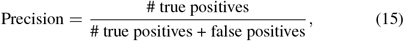

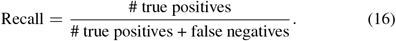

#### 2.3.3 Real-world dataset

To assess the performance of STIR on real data, we analyzed 78 major depressive disorder (MDD) subjects and 79 healthy controls (HC) from the RNA-Seq gene network module analysis in Ref. (Le *et al*. (ress)). This dataset consists of whole blood RNA-Seq measurements of 5,912 genes for each subject. Sequencing yielded an average of 30 million reads per individual, and gene expression levels were quantified from reads of 19,968 annotated protein-coding genes, followed by low read-count and outlier removal as well as technical and batch effect adjustment. Coefficient of variation filtering resulted in the final set of 5,912 genes that we use in the current study to test for association with MDD status (Le *et al*. (ress)).

## 3 Results

### 3.1 Comparison of the performance of STIR with Relief-based permutation

Our first aim is to determine whether the more computationally efficient pseudo t-test approach of STIR is a reliable alternative to a model-free permutation test. We use multiSURF as the neighborhood algorithm in STIR, but constant *k* algorithms are expected to perform similarly (see following subsection). We also use the multiSURF neighborhood for permutation-Relief. Using an FDR adjusted p-value threshold *α* = 0.05, we observe that STIR (mauve) and permutation-Relief (blue) indeed perform nearly the same in both main effect and interaction effect simulations in terms of True Negatives, Precision, and Recall (Fig. 2). For completeness and to provide an indicator of power, we also compare STIR with the performance of a univariate t-test (green). For main effect simulations (Fig. 2A), all methods have a similarly low Recall because the simulated main effect size and sample size were chosen to be relatively low and challenging.

**Fig. 2.**
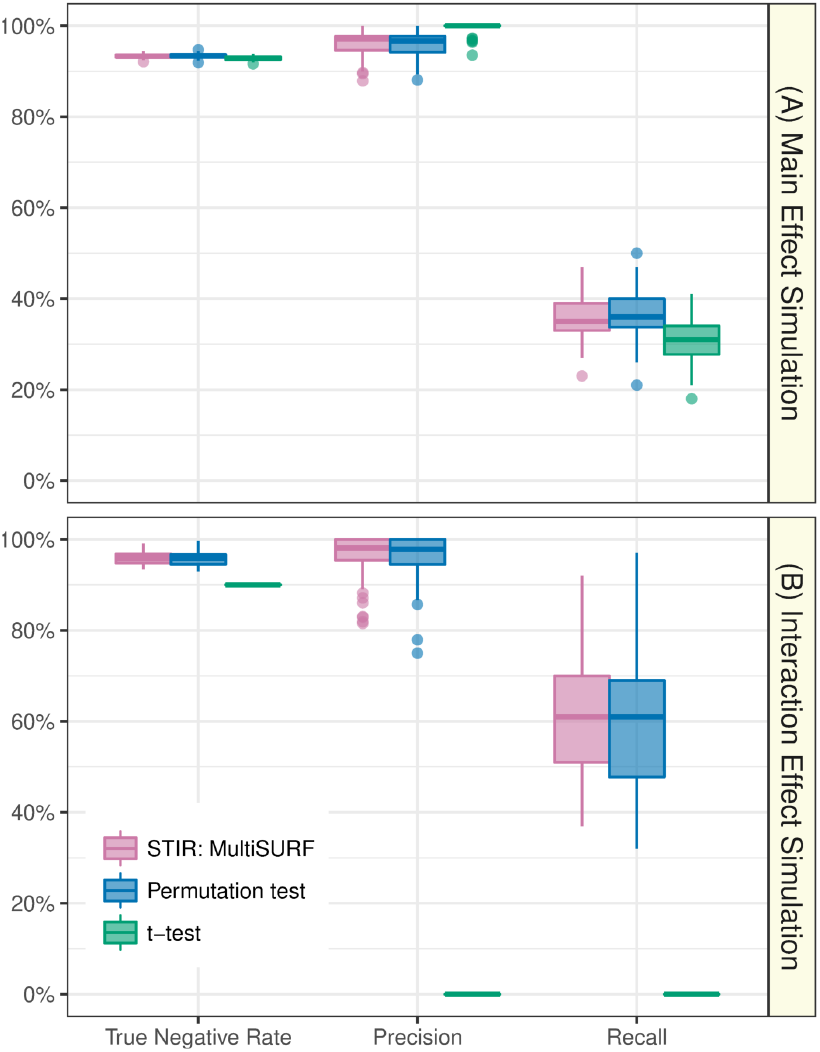
STIR versus permutation-test multiSURF and univariate t-test. Comparison of the performance (True Negative Rate, Precision, and Recall) of STIR (with multiSURF neighborhood, mauve), permutation test of multiSURF (blue), and univariate t-test (green) to detect functional attributes. Each method determines positives by 0.05 FDR adjusted p-value threshold. Each simulation is replicated 100 times with *m* =100 samples and *p* = 1000 attributes with 100 functional (A) main effects (bias=0.8) and (B) interaction network effects (*s*_int_ = 0.4).

As we discuss more in Sec. 3.2 (Fig. 3), for main effects, it is possible to further increase the Recall of STIR beyond a univariate t-test if one uses STIR with ReliefF and a larger *k* (up to the maximum *k*_max_ = ⌊(*m* — 1)/2⌋); however, this *k* would cause a decrease in performance for interactions relative to STIR with lower values of *k*. The multiSURF neighborhood constitutes a compromise between main effect and interaction effect performance, as we explore more below.

**Fig. 3.**
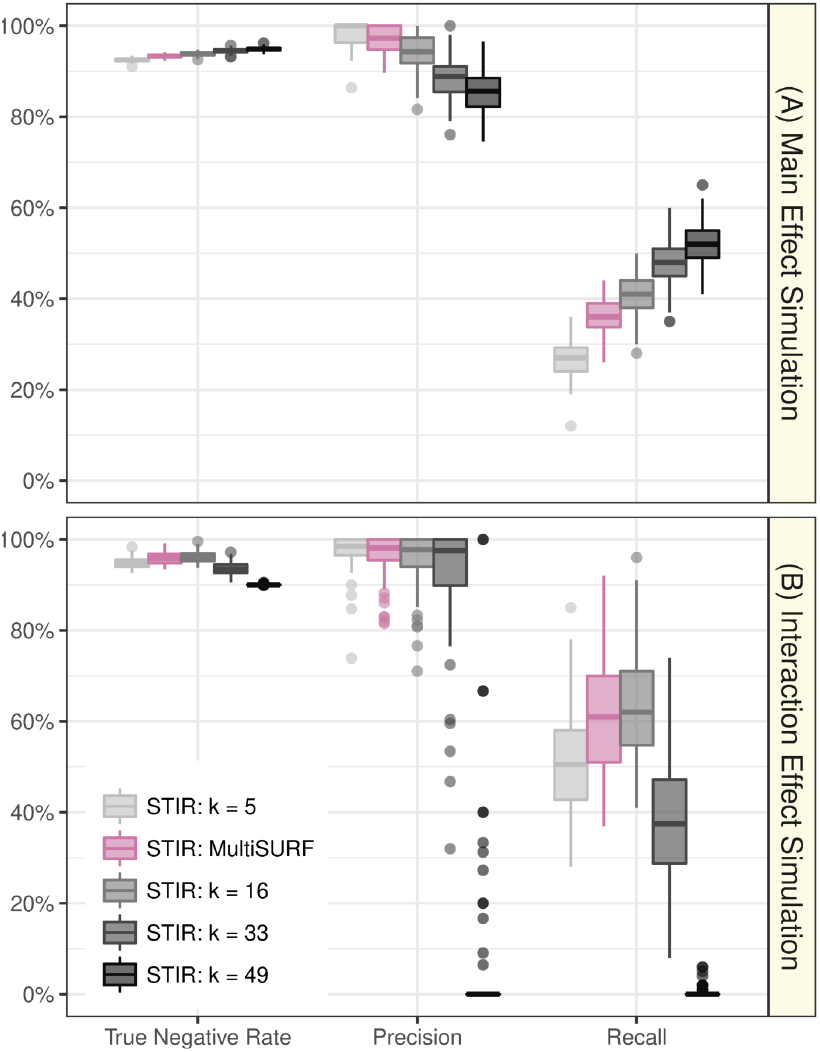
The effect of *k* on the performance of STIR to detect functional attributes with main effects (A) and interaction effects (B). Comparison of the performance (True Negative Rate, Precision, and Recall) of STIR-ReliefF for multiple values of nearest neighbors *k* (*k* = 5, 16, 33, 49, gray scale) and STIR-multiSURF (adaptive radius, mauve). All methods determine positives using a 0.05 FDR adjusted p-value threshold. Each simulation is replicated 100 times with *m* = 100 samples and *p* = 1000 attributes with 100 functional.

For interaction simulations (Fig. 2B), the t-test still has a similarly high True Negative Rate to STIR. However, this high rate is because no t-tests are true positive: there are no main effects and the t-test has zero Precision and Recall. STIR on the other hand still has high Precision and Recall (Fig. 2B) because Relief-based methods are sensitive to interactions among attributes (provided the number of neighbors is not too large).

For a dataset of the size simulated in our study (*m* = 100 samples and *p* = 1, 000 attributes) STIR has a2.1-second runtime on adesktop with an Intel Xeon W-2104 CPU and 32GB of RAM. As expected, a permutation test with 10,000 replicates takes approximately 10,000 times longer: over 5 hours on the same desktop. Thus, STIR provides a significant time savings over permutation for computing p-values.

Our next aim is to gain insight into the performance of STIR with a ReliefF neighborhood (fixed *k* neighbors) and how its performance relates to STIR with a multiSURF neighborhood (adaptive radius). In the main effect simulations (Fig. 3A), as *k* increases, STIR gains more power to detect the functional attributes (increasing Recall) and with an expected increase in false positive attributes (decreasing Precision). The increasing Recall with *k* is expected for main effects because ReliefF becomes more myopic (more like a univariate t-test) as *k* increases (Robnik-Šikonja and Kononenko (2003); McKinney *et al*. (2013)). The increase in Recall is limited in part by the maximum number of neighbors being *k*_max_ = |⌊(*m* — 1)/2⌋ = 49.

### 3.2 The effect of *k* in detecting functional attributes

In contrast, for interaction simulations (Fig. 3B), the relationship between *k* and Recall is no longer monotonic. Rather, the Recall reaches a maximum at approximately *k* = *m*/6 and this performance is similar to using the adaptive radius in multiSURF. As *k* increases beyond *k* = *m*/6 to the maximum *k*_max_, ReliefF becomes more myopic and has nearly zero Precision and Recall. This result corroborates the findings in Ref. Urbanowicz *et al*. (2018a) that multiSURF is sensitive to two or threeway interactions. However, we also note that the STIR-ReliefF with *k* = |⌊*m*/6⌋ = 16 results are similar to STIR-multiSURF for main effect and interaction effect simulations (because the average *k* in multiSURF is close to *m*/6). These versions of STIR will yield similar results for balanced data that are optimal for detecting interactions while being reasonably powerful for main effects. STIR_*k*=*m*/6_ has a computational speed advantage over STIR-multiSURF, but STIR-multiSURF may have an advantage when there is class imbalance (Urbanowicz *et al*. (2018 a)). If one wanted to optimize the sensitivity of STIR for main effects and neglect interactions, one would use STIR_*k*=*k*_max__. Furthermore, in all simulation scenarios, the correlation between STIR scores (pseudo t-statistic) and the original Relief-based scores (diff function) are above 0.98 (see Supplement Fig. 1 for more detail).

### 3.3 Real-world data

We apply STIR to the RNA-Seq study of 78 major depressive disorder (MDD) subjects and 79 healthy controls described in Ref. (Le *et al*. (ress)). The dataset contains 5,912 genes after preprocessing and filtering (see Methods for more detail). Using an FDR threshold of 0.05, STIR with the multiSURF neighbor-finding method detects 32 statistically significant associations **(mauve** and **gray** genes above the dashed horizontal line in Fig. 4). These 32 significant STIR genes include all eight of the genes that passed the 0.05 FDR threshold from the standard t-test method **(gray** genes to the right of the vertical dashed line in Fig. 4). Thus, in addition to its documented ability to identify interactions, STIR also has high power to detect main effects among its FDR-adjusted significant genes.

**Fig. 4.**
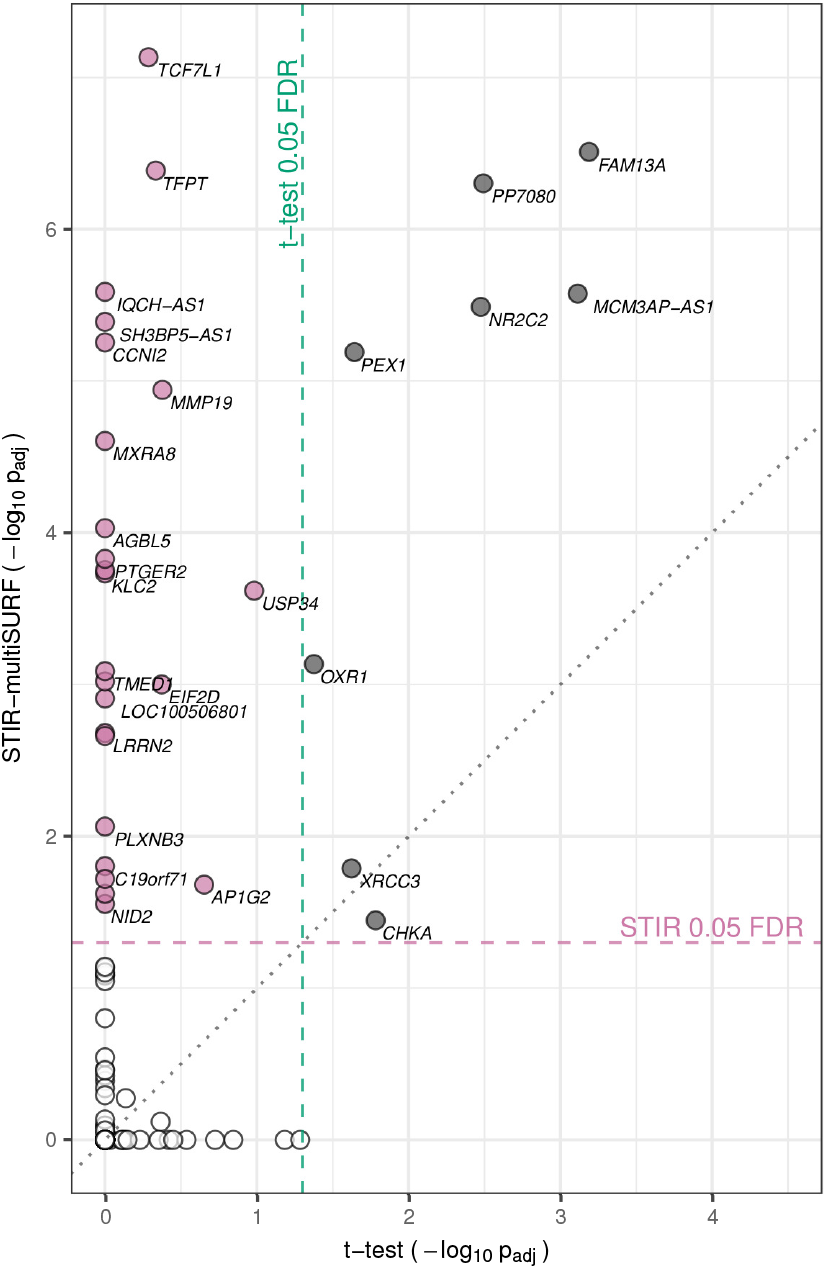
Major depressive disorder gene scatter plot of — log_10_ adjusted significance for STIR-multiSURF and standard t-test for RNA-Seq differential expression. STIR-multiSURF finds 32 genes that are significant at the FDR-adjusted 0.05 level (above horizontal dashed line). Standard t-test finds eight genes that are significant at the FDR-adjusted 0.05 level (to right of vertical dashed line). STIR identifies all eight significant main effects from the t-test (gray) and additional candidate genes (mauve) that may involve interactions. Due to overlap of plot points, not all significant genes are labeled. See Supplementary Fig. S2 for detailed labels.

The STIR-multiSURF genes that are outside of the intersection with the t-test (mauve) such as TCF7L1, a component of the Wnt signaling pathway, may be good candidates for interaction effects. An extended Venn diagram of the gene-significance overlap of these two methods is provided in Supplementary Fig. S2. Although beyond the scope of the current study, characterization of interactions could be performed to create an expression-epistasis network from the STIR MDD genes (McKinney *et al*. (2009); Lareau *et al*. (2015)) and help identify underlying mechanisms of MDD susceptibility.

Using STIR with fixed *k* = *m*/6, we identified 41 FDR-adjusted significant genes (not plotted). These 41 STIR *k* = *m*/6 genes include 31 of the 32 STIR-multiSURF genes. Thus, as we found in the simulation studies, these two versions of STIR perform similarly, with multiSURF being more conservative. Future studies will address the replication of the statistically significant STIR effects, the characterization of STIR interactions and the mathematical connection between neighbor-finding methods in STIR. The STIR runtime for the RNA-Seq data was approximately 19 seconds on a desktop with an Intel Xeon W-2104 CPU and 32GB of RAM.

## 4 Discussion

To our knowledge, STIR is the first method to use a theoretical distribution to calculate the statistical significance of Relief attribute scores without the computational expense of permutation. Previously, it was difficult to assess the false discovery rate of Relief-based attribute lists because arbitrary thresholds were used. STIR is able to report statistical significance of Relief-based scores by a pseudo t-test that accounts for variance in the mean difference of miss and hit nearest neighbor diffs. We assessed STIR’s power and ability to control false positives using realistic simulations with main effects and network interactions. We applied STIR to real data to demonstrate the identification of biologically relevant genes.

We showed that the statistical performance using STIR p-values is the same as using permutation p-values. This validates the STIR pseudo t-test and means one can use it instead of costly permutation testing. We chose the number of permutation to be 10,000 to minimize the computational expense while obtaining accurate permutation p-values. Specifically, if only 1,000 permutations were performed, the p-values would be bounded below by 0.001, which would lead to an inflation of insignificant tests after FDR correction (*p_adj_* > 0.05) in simulated datasets with 1,000 attributes. Nevertheless, 10,000 permutations requires considerable computation time, especially in large datasets such as the analyzed gene expression data. Hence, by showing very similar performance to permutation, STIR shows an efficient implementation to compute the p-value for each attribute while producing scores that are highly correlated with the standard Relief-based scores.

We showed the STIR formalism generalizes to all Relief-based neighbor finding algorithms, including MultiSURF. We showed that STIR-MultiSURF and STIR_*k*=*m*/6_ perform similarly for main effect and interaction simulations. This suggests that one may prefer to use constant-*k* STIR_*k*=*m*/6_ for the computational speed advantage; however, we have not tested the statistical performance for imbalanced data. Our results suggest that power for detecting interactions is maximized near *k* = *m*/6 (higher or lower *k* decreases the power). Power for detecting main effects is highest with the myopic maximum *k* = *k_max_* = ⌊(*m* — 1)/2⌋. Real biological data will likely contain a mixture of main effects and epistasis network effects (McKinney and Pajewski (2012)). The value *k* = *m*/6 is a good compromise because it maximizes the radius for detecting interactions while still giving reasonable power for detecting main effects. However, the STIR formalism may help tune the elements of an attribute-specific *k* vector, where each attribute, *a*, is allowed to use a different *k_a_* to preferentially detect a main effect or interaction effect as informed by the data (McKinney *et al*. (2013)). For those using a constant-*k* (ReliefF) approach, our results suggest that using *k* = *m*/6 may offer a better default than the pervasive use of *k* = 10, which was an arbitrary choice in the early literature.

Our simulation study focused on obtaining a quality assessment of statistically significant STIR associations between an attribute and the outcome while taking into account the complex underlying architecture of interactions among attributes. Therefore, the simulation is designed to generate realistic and challenging datasets leading to relatively low Recall. In datasets with larger sample size (*m* = 200), we observe higher Recall values but otherwise similar findings as presented in the Results section (results for *m* = 200 not shown). Furthermore, from a machine learning point of view, if the researcher wishes to include more attributes in their subsequent analysis, they may increase the FDR threshold to allow for more false positives and improve the Recall value. A future study that analyzes this Recall/Precision trade-off would prove valuable in understanding statistical characteristics of selected features from Relief-based methods.

The STIR score improves the standard Relief-based scores because, rather than simply being a difference of means, STIR incorporates within and between group variances. Moreover, this pseudo t-test score can be transformed into a p-value. The advancement of STIR over Relief-based scores is similar to going from a fold change to describe differential expression to a t-test. The assumptions of a t-test-independent observations and normality of the population distributions – are not satisfied for the STIR test in general, which is why we refer to it as a pseudo t-test. When the average number of neighbors *k* is sufficiently large, duplicate pairs will occur in the estimate of the average hit and miss diffs. The dependence induced by duplicate neighbors may increase the false positive rate because the variance estimates are narrowed, the STIR statistics inflated, and the p-values deflated. One could simply remove duplicates; however, the duplicates are beneficial with respect to power because they add weight to pairs of instances that are very similar to each other. The effect of duplicates has a similar effect as a distance-based weighting scheme such as the exponential decaying influence of neighboring instances used in some Relief-based algorithms (Robnik-Šikonja and Kononenko (2003)).

A related approach to reduce the dependence-induced false positive rate is to perform sub-sampling of the neighbor pairs, which reduces duplicates but maintains some distance-based weighting. An alternative approach would be to incorporate variance regularization into the STIR statistic to inflate the variance to a level consistent with independent neighbors. Despite the dependence of neighbors, our empirical results show that, even when unmodified, the STIR pseudo t-test shows comparable performance with permutation test in both simulation scenarios with main and interaction effects.

Transformations such as the square root help increase the normality of the distribution of distances. However, to stay close to the original Relief score formula, we did not transform the distance values in the results shown here, but the transformation is provided as an option via the `transform` parameter of the STIR function in our software. Preliminary analysis indicates little difference when transformation is applied (results not shown).

It has been shown that Relief-based algorithms benefit from the iterative removal of the worst attributes and then repeating the estimation of the remaining attributes. Thus, another future direction is to develop a strategy for STIR that incorporates iterative attribute removal in a way that minimizes the false positives due to iteration-induced multiple testing. STIR feature selection could be embedded in the backwards elimination of pEC for feature selection and classification (Le *et al*. (2017)) or embedded in a nested cross-validation approach. Effective strategies also must be developed for testing for replication of significant STIR effects because typical replications do not have dependence among other features, whereas Relief scores depend on the context of other variables in the data.

Extensions of STIR will involve multi-class data, quantitative trait data (regression) and correction for covariates. Just as an ANOVA extends the t-test to multiple conditions, we anticipate the extension of STIR to multi-state will involve an ANOVA formalism and F-test. Similarly, we envision regression-STIR to follow a linear model formalism. The current implementation of STIR does not deal with missing data. In a future implementation, to handle missing data we will modify the diff to estimate the probability that two instances (one or both possibly missing) have different values conditioned on their class. Application to GWAS data requires no additional modifications other than specification of a different diff function for categorical variables. Future studies will apply STIR to GWAS as well as eQTL and other high dimensional data to identify interaction effects.

## Acknowledgements

**Funding**

This work was supported in part by the National Institute of Health Grant Nos. LM010098 and LM012601(to JHM) and Nos. GM121312 and GM103456 (to BAM).

## References

Benjamini, Y., Drai, D., Elmer, G., Kafkafi, N., and Golani, I. (2001). Controlling the false discovery rate in behavior genetics research. Behavioural brain research, 125(1-2), 279–284.

Greene, C. S., Penrod, N. M., Kiralis, J., and Moore, J. H. (2009). Spatially Uniform ReliefF (SURF) for computationally-efficient filtering of gene-gene interactions. BioData Mining, 2, 5.

Kira, K. and Rendell, L. A. (1992). The feature selection problem: Traditional methods and a new algorithm. In Proceedings Tenth National Conference on Artificial Intelligence, pages 129–134. AAAI Press/The MIT Press.

Kononenko, I., Šimec, E., and Robnik-Šikonja, M. (1997). Overcoming the Myopia of Inductive Learning Algorithms with RELIEFF. Applied Intelligence, 7(1), 39–55.

Lareau, C. A., White, B. C., Oberg, A. L., and McKinney, B. A. (2015). Differential co-expression network centrality and machine learning feature selection for identifying susceptibility hubs in networks with scale-free structure. BioData mining, 8(1), 5.

Le, T. T., Simmons, W. K., Misaki, M., Bodurka, J., White, B. C., Savitz, J., and McKinney, B. A. (2017). Differential privacy-based evaporative cooling feature selection and classification with relief-f and random forests. Bioinformatics, 33(18), 2906–2913.

Le, T. T., Savitz, J., Suzuki, H., Misaki, M., Teague, T. K., White, B. C., Marino, J. H.,Wiley, G., Gaffney, P. M., Drevets, W. C., McKinney, B. A., Bodurka, J., and McKinney, B. A. (in press). Identification and replication of rna-seq gene network modules associated with depression severity. Translational Psychiatry.

McKinney, B. and Pajewski, N. (2012). Six degrees of epistasis: statistical network models for gwas. Frontiers in genetics, 2, 109.

McKinney, B. A., Crowe, J. E., Guo, J., and Tian, D. (2009). Capturing the spectrum of interaction effects in genetic association studies by simulated evaporative cooling network analysis. PLoS genetics, 5(3), e1000432.

McKinney, B. A., White, B. C., Grill, D. E., Li, P.W., Kennedy, R. B., Poland, G. A., and Oberg, A. L. (2013). ReliefSeq: A Gene-Wise Adaptive-K Nearest-Neighbor Feature Selection Tool for Finding Gene-Gene Interactions and Main Effects in mRNA-Seq Gene Expression Data. PLOS ONE, 8(12), e81527.

Park, S. and Lehner, B. (2013). Epigenetic epistatic interactions constrain the evolution of gene expression. Molecular systems biology, 9(1), 645.

Robnik-Šikonja, M. and Kononenko, I. (2003). Theoretical and empirical analysis of relieff and rrelieff. Machine learning, 53(1-2), 23–69.

Urbanowicz, R. J., Olson, R. S., Schmitt, P., Meeker, M., and Moore, J. H. (2018a). Benchmarking relief-based feature selection methods for bioinformatics data mining. Journal of Biomedical Informatics.

Urbanowicz, R. J., Meeker, M., Cava, W. L., Olson, R. S., and Moore, J. H. (2018b). Relief-based feature selection: Introduction and review. Journal of Biomedical Informatics.

